# Effects of abiotic and biotic drivers on tadpoles in seasonal rock pools of Western Ghats rock outcrops, India

**DOI:** 10.1101/2024.04.13.589092

**Authors:** Vijayan Jithin, Rohit Naniwadekar

**Author notes:** Correspondence: Rohit Naniwadekar, Nature Conservation Foundation, Mysore, Karnataka, India.;, Vijayan Jithin, Nature Conservation Foundation, Mysore, Karnataka, India.

## Abstract

We assessed the influence of abiotic (pool size, monsoon progression) and biotic (predator abundances) factors on occurrence and abundance of three species of tadpoles by periodically monitoring rock pools in lateritic plateaus. Tadpole responses were positively associated with rock pool size and monsoon progression negatively, but not with predator abundance.

## 1. INTRODUCTION

Frogs have biphasic life histories, and most spend a significant amount of time in their larval form (tadpoles), a stage that can have implications for their adult populations (Berven, 1990; Liedtke et al., 2022). Despite this, past research has mostly focused on studying adult frogs (Vera Candioti et al., 2023; Annibale et al., 2023). Tadpoles’ survival depends on various biotic and abiotic factors in the aquatic habitat, including predation, competition, aquatic permanence, temperature and anthropogenic disturbances, among others (Van Buskirk & Smith, 2021; Alford, 1999; Cayuela et al., 2020).

Freshwater rock pools (FRPs) formed due to erosion and weathering in outcrops are critical frog breeding habitats (Jocque et al., 2010). They vary in abiotic features like size and hydroperiod, and biotic features like predator density and competition, which are known to directly or indirectly influence tadpole occurrence (Aristizábal Botero, 2023; Anusa et al., 2012; Buxton & Sperry, 2017). These temporally dynamic habitats varying in biotic and abiotic features facilitate quantification of population- and community-structuring processes that are difficult to quantify in larger, complex systems (Brendonck et al., 2010).

Lateritic rock outcrops in the northern part of the Western Ghats-Sri Lanka Biodiversity hotspot are abundant with FRPs, observed as depressions on the rocky substrata with pan- or bucket-shaped geomorphological structures, fed by monsoon rains (Figure 1b). They harbor various endemic organisms uniquely adapted to the highly variable environment in the open ecosystem (Watve, 2013; Gaitonde et al., 2016). There is existing information on arthropod and aquatic plant communities (Sheth et al., 2019; Kulkarni et al., 2022; Kulkarni et al., 2023), but vertebrate ecology, especially of tadpoles, is lacking. Jithin *et al*. (2023) showed that large FRPs are important for adults of three species of frogs: *Euphlyctis jaladhara, Microhyla nilphamariensis*, and *Polypedates maculatus*; and recommended restoration or creation of rock pools for amphibian conservation in light of reduced FRP availability because of rapid conversion of outcrops to orchards that reduces the availability of FRPs. Knowledge of tadpole ecology in the FRPs is essential for further research and conservation actions.

**Figure 1.**
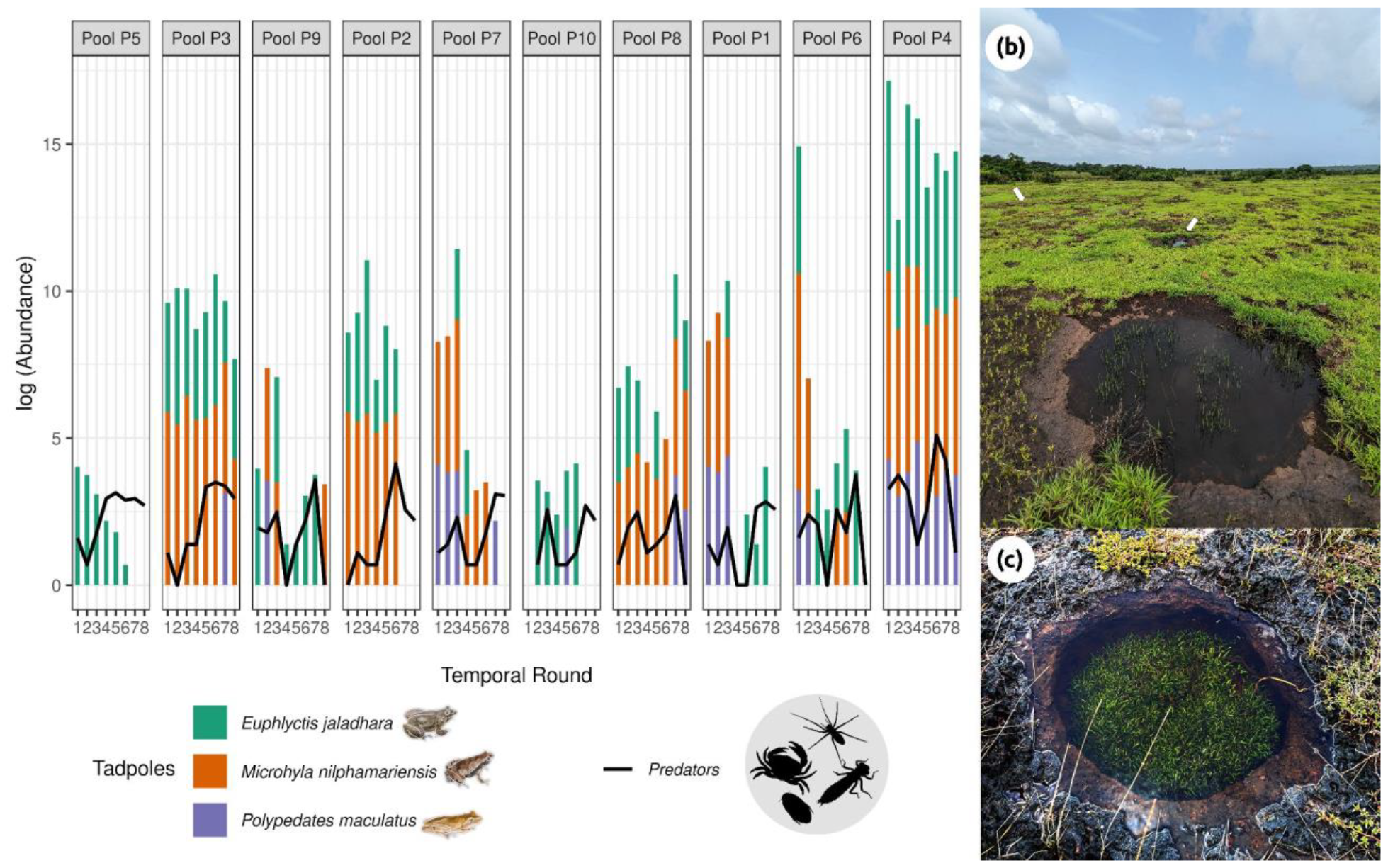
(a) Barplots showing the abundance of tadpoles counted in each pool across the temporal rounds along monsoon progression. Black lines indicate the overall abundance of predators. The pools are ordered from smallest to largest pool volume. (b) Photograph showing the rockpool distribution in the rock outcrop landscape of the northern Western Ghats, white pointers indicate positions of other rock pools; (c) an individual rock pool in closeup. Photographs and illustrations by Vijayan Jithin.

We investigated the influence of FRP size, predator abundances and monsoon progression on the occurrence and abundance of tadpoles of three species of anurans: Jaladhara Skittering Frog (*Euphlyctis jaladhara*), Nilphamari Narrow-mouthed Frog (*Microhyla nilphamariensis*), and Common Indian Treefrog (*Polypedates maculatus*). We determined the relative influence of abiotic (pool size and monsoon progression) and biotic (predator abundance) factors on the occurrence and abundance of three species of tadpoles, as pond permanence and predation are known as powerful forces shaping the structure of tadpole communities (Skelly, 1997). We expected that tadpole occurrence and abundance would be 1) positively associated with pool size since larger pools will hold water during non-rainy days, thereby protecting tadpoles from desiccation, 2) negatively associated with monsoon progression as the end of monsoon poses survival risks due to desiccation, and 3) negatively associated with the abundance of potential predators.

## 2. METHODS

VJ monitored 10 FRPs spread over an area of 1.15 km^2^ in the low-elevation, privately-owned lateritic rock outcrop (151–190 m a.s.l.) of Devihasol in Ratnagiri District of Maharashtra state in west India (16°44’–16°45’N; 73°25–73°27’E; Figure 1 c-d). The area receives the south-west monsoon rainfall from June to September, with sporadic rainfall in May and October. We sampled pools between July and September 2022. The monthly rainfall during the study period varied from 408.1 mm in September to 639.8 mm (July) in the study area, with the number of rainy days ranging from 18-23 (data from Nate, Rajapur; Department of Agriculture, Government of Maharashtra, 2024). VJ conducted nighttime pool surveys for tadpoles (Scott Jr. & Woodward, 1994). All 10 FRPs were monitored eight times during the study period between 1900–2200 h, usually in clear weather, barring occasional rain incidences. The pool water was clear during all the observation occasions.

The observer gently walked along the bank and scanned the pool to record all animals (Scott Jr. & Woodward, 1994). Care was taken not to recount the same schools of tadpoles, and a red light was used while approaching the pool to avoid light disturbance, following Jithin *et al*. (2022). The observer enumerated tadpoles of the three species and their potential invertebrate predators: fishing spiders (*Araneae*), crabs (*Decapoda*), dragonfly larvae (*Odonata*), and water beetles (*Coleoptera*) (Skelly, 1997; Valdez, 2020; Mogali et al., 2022), by counting them using head and hand-held torch lights. Pool volume (cm^3^) was calculated by multiplying the mean depth, maximum length (longest axis) and maximum width (longest axis perpendicular to length). The pool water volumes were calculated four times during the study period, and the largest value was considered the maximum pool volume, which was log-transformed for analysis. We used log-transformed values of predator abundances that were estimated during each temporal replicate. We confirmed the identity of tadpoles using published literature on tadpoles (Raj et al., 2023; Altig & McDiarmid, 1999).

We performed all the analyses in R (v. 4.3.0) (R Core Team, 2024). We used the Hierarchical Modelling of Species Communities framework (HMSC) (Ovaskainen et al., 2017), belonging to the class of joint species distribution models, to understand how each species responded to the predictors. We used the temporal replicate ID, and the spatial arrangement of pools (x-y coordinates) as random levels in the model. Our dataset consisted of 79 temporal replicates across 10 pools. Our predictor variables were maximum pool volume, monsoon progression phase (July, August, September), and predator abundances (fishing spiders, crabs, dragonfly larvae, water beetles). We applied a hurdle model approach to model the occurrence (presence-absence) with probit regression, and log-transformed abundances conditional on presence with linear regression using ‘Hmsc’ package, assuming the default prior distributions (Tikhonov et al., 2020; Ovaskainen and Abrego, 2020). Posterior sampling for three Markov Chain Monte Carlo (MCMC) chains was performed, each of which we sampled for 300,000 iterations, out of which the first 50,000 were discarded as a transient, and thinned the remainder by 1,000 to yield 250 posterior samples per chain. Before fitting the final model, we explored the MCMC convergence of models with thin values of 1, 10, and 100. We assessed the convergence of the model using Gelman diagnostics and evaluated the explanatory power of the model using Tjur’s *R*^*2*^ for the presence-absence model and normal *R*^*2*^ for the model with scaled log abundance, conditional on presence. Relative contributions of fixed and random effects in explaining the variation in species occurrence and abundances, was assessed using the variance partitioning approach. We evaluated the influence of predictors on species occurrence and abundance by examining the ≥ 95% posterior probability.

## 3. RESULTS

The pool volumes ranged between 0.04 and 7.46 m^3^ and the mean depth ranged between 0.1 and 0.27 m. While we encountered *Euphlyctis* tadpoles in all the pools, *Microhyla* and *Polypedates* occurred in eight of the 10 pools (Figure 1).

Measured by *R*^*2*^, our fitted HMSC model explained, on average, 36% of the variation in species occurrence, and 53% in species abundance (conditional on presence). The occurrence of all tadpole species was negatively associated with monsoon progression (explained variation: 22.5%), and positively associated with pool volume (explained variation: 26.8%) for *Polypedates* and *Microhyla* (Figure 2). The abundance (conditional on presence) was positively associated with pool volume (explained variation: 25.7%) for *Euphylctis* and *Microhyla*, and negatively associated with monsoon progression (explained variation: 26.1%) for all tadpole species (Figure 2). Predator abundance explained less than 7% of the variation for all potential predators and did not show statistically significant associations. We did not find any statistically significant support for spatial (α_1_ = 0) and temporal signal (α_2_ = 0) in the residuals, indicating a lack of spatial and temporal autocorrelation (Figure 2a).

**Figure 2.**
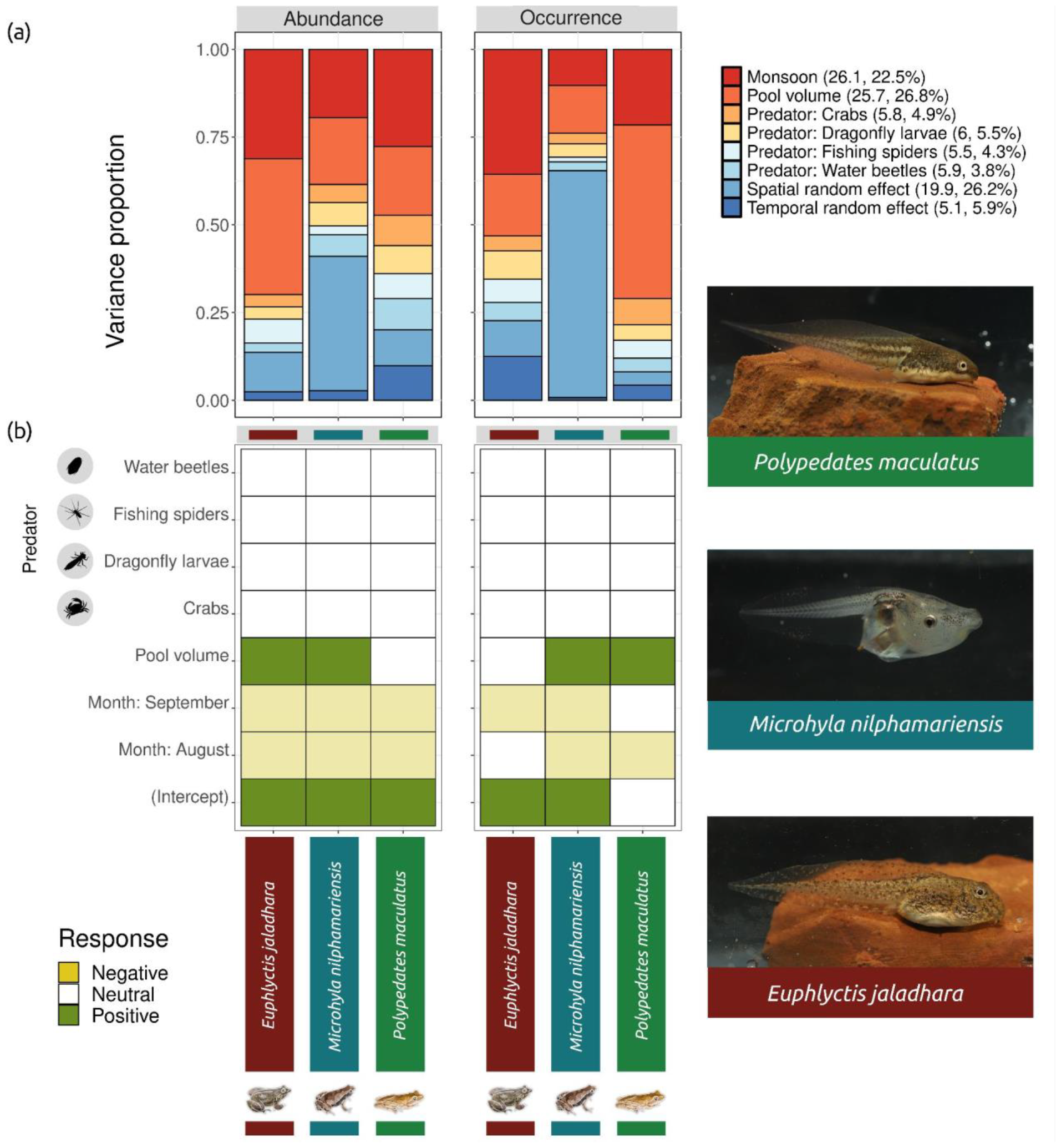
HMSC model results showing (a) the variance partitioning of explained variation among environmental covariates and random effects, with values in parentheses of legend markers, show the mean values across species for conditional abundance and occurrence respectively; (b) the mean posterior regression (*β)* parameter values measuring the species-specific responses of tadpoles to each of the environmental covariates. Yellow color indicates negative responses and olive green color positive responses with ≥ 0.95 posterior probability. The tadpole photographs show individuals at the developmental stage of (i) *Polypedates maculatus* - 37; (ii) *Microhyla nilphamariensis* - 28; and (iii) *Euphlyctis jaladhara* - 31 (Gosner, 1960). Photographs by Rohit Naniwadekar. Illustrations by Vijayan Jithin.

## 4. DISCUSSION

Given our expectations, we found support for the role of abiotic factors, i.e. monsoon progression and pool volume, in influencing tadpole occurrence and abundance in the FRP of the lateritic outcrops in the northern Western Ghats. However, we did not find support for the role of predators. This fills a valuable knowledge gap on a critical life history stage of multiple peninsular Indian endemic amphibians and its association with abiotic factors. Previous studies from the Western Ghats have shown that abiotic factors such as stream slope, water depth, temperature, canopy cover and water velocity influence endemic tadpole distribution in the streams (Priti et al., 2021; Thomas et al., 2019; Girish & Krishnamurthy, 2009). However, they have not explicitly tested for the influence of biotic factors like predators. This limited information from forest streams is complemented by our insights from rock outcrop pools, which are highly vulnerable to extreme climatic fluctuations.

Monsoon progression significantly influenced the occurrence and abundance of tadpoles in rock pools. End of monsoon would increase the risk of desiccation for the growing tadpoles. Incidentally, we found lower incidences of *Microhyla* and *Polypedates* adults towards the end of monsoon (Jithin et al., 2023). This indicates lowered breeding intensity towards the end of monsoon, which may also contribute to lower tadpole occurrence and abundance.

Pool volume influenced tadpole occurrence and abundance. The association between larger pool volume with higher tadpole abundance and occurrence of some species in our study is consistent with insights from rock pools in the Atlantic Forests of Brazil (Schiesari et al., 2018). This is possibly through the influence of pool volume on the hydroperiod (Brooks & Hayashi, 2002). Pool volume and depth are widely known as the best predictors of animal occurrence in freshwater pools, with large and deep pools having high food availability and fewer predators, resulting in greater reproductive success of amphibians (Rothenberger & Baranovic, 2021; Rothenberger et al., 2019).

We did not find statistically significant support for the role of predator abundance. Importantly, predation by permanent aquatic predators such as fish is considered the most important biotic factor shaping temporal and spatial composition of tropical tadpole communities (Heyer et al., 1975). The ephemeral rock pools we observed were mostly devoid of fish and mainly harbored invertebrate tadpole predators. These findings are in line with the predictions of Heyer et al., (1975), who expected biological factors such as predation to play a lesser role than the abiotic factor of pools drying up, in the reproductive success of opportunistically breeding frogs in small, ephemeral habitats. In the context of rock pool habitats, more natural history information on the temporal growth and development patterns of both the amphibian and predator species, and experimental approaches will better elucidate the role of biotic factors such as predation and interspecific competition (Heyer et al., 1975).

Given the differing responses of tadpoles to different predictors, as revealed by the variance partitioning analysis, our results also highlight the importance of species-specific tadpole ecology studies, which can inform species-specific conservation strategies. Considering the global threats amphibians are facing from habitat degradation and land conversion, and particularly agricultural land-use change in the rock outcrops (Luedtke et al., 2023; Jithin et al., 2023), conservation strategies need to be developed based on both adult and tadpole ecology in the local context. Our study provides the first insights for rock pool restoration in the northern Western Ghats rock outcrops, based on tadpole ecology, complementing the previous work on adult frogs.

## AUTHOR CONTRIBUTIONS

RN and VJ conceived the ideas and designed the methodology. VJ collected the data and analyzed it with input from RN. VJ led the writing of the manuscript with inputs from RN. All authors contributed critically to the drafts and gave final approval for publication.

## ACKNOWLEDGEMENTS

We thank the Maharashtra Forest Department, particularly the Chief Wildlife Warden, Sunil Limaye, for giving us the necessary permits (Letter No. Desk-22(8)/WL/Research/CR-53(20-21) /3361/22-22) to conduct the study. We thank On the Edge (UK), The Bombay Environmental Action Group and The Habitat Trust (India) for funding this work. We thank Pooja Pawar, Jahnavi Joshi, Aparna Watve, Varad B. Giri, Manali Rane, and Arpitha Jayanth for useful discussions and support during the study; Aditya Gadkari, Ajay Nachinekar, Chandrakanth Gurav, Harshad Tulpule, Kamalakar Gurav, Pooja Ghate, Poorva Joshi, Pradeep Dingankar, Rakesh Patil, Rashmi Karandikar, Ravindra Karandikar, Santosh Padhye, Shailesh Joshi, Suhas Gurjar, Sujan Dandekar, and Yash Vichare for support during fieldwork. RN thanks Anand Osuri, Kulbhushansingh Suryawanshi, Suri Venkatachalam, and Madhura Niphadkar for useful discussions.

## ETHICAL CONSIDERATIONS

We did not collect samples of, or preserve animals as part of this observational study. We have obtained permission to study amphibians in the study area from the Principal Chief Conservator of Forests (Wildlife), Maharashtra State, India (Letter No. Desk-22(8)/WL/Research/CR-53 (20-21)/3361/22-22) and have obtained necessary institutional ethics committee approval from the Nature Conservation Foundation-India.

## CONFLICT OF INTEREST STATEMENT

The authors declare no conflict of interest.

### DATA AVAILABILITY STATEMENT

Data and codes used in this study will be uploaded to DataDryad upon acceptance.

## Notes

### Competing Interest Statement

The authors have declared no competing interest.

## REFERENCES

1. Alford, R. A. (1999). Ecology: Resource use, competition, and predation. In: McDiarmid, R. W. and R. Altig (Eds.). Tadpoles: The Biology of anuran larvae (pp. 240–278). The University of Chicago Press.

2. Altig, R., & McDiarmid, R. W. (1999). Diversity: Familial and generic characterizations. In: McDiarmid, R. W. and R. Altig (Eds.). Tadpoles: The Biology of anuran larvae (pp. 295–337). The University of Chicago Press.

3. Annibale, F. S., Wassersug, R. J., Rossa-Feres, D. D. C., Nomura, F., Brasileiro, C. A., Sabbag, A. F., … & Phillips, J. R. (2023). The case for studying tadpole autecology, with comments on strategies to study other small, fast-moving animals in nature. Austral Ecology, 48(5), 855–876. 10.1111/aec.13367

4. Anusa, A., Ndagurwa, H. G. T., & Magadza, C. H. D. (2012). The influence of pool size on species diversity and water chemistry in temporary rock pools on Domboshawa Mountain, northern Zimbabwe. African Journal of Aquatic Science, 37(1), 89–99. 10.2989/16085914.2012.666378

5. Aristizábal Botero, Á. (2023). Ecological dynamics of macroinvertebrates from inselbergs of the Colombian Guiana Shield. PhD Thesis. Aquatic Zoology and Ecology Laboratory, Department of Biology, Universidad de Los Andes, Bogotá, Colombia, and the Community Ecology Laboratory, Department of Biology, Vrije Universiteit Brussel, Brussels, Belgium.

6. Berven, K. A. (1990). Factors affecting population fluctuations in larval and adult stages of the wood frog (Rana sylvatica). Ecology, 71(4), 1599–1608. 10.2307/1938295

7. Brendonck, L., Jocque, M., Hulsmans, A., & Vanschoenwinkel, B. (2010). Pools” on the rocks”: freshwater rock pools as model system in ecological and evolutionary research. Limnetica, 29(1), 0025–40. 10.23818/limn.29.03

8. Brooks, R. T., & Hayashi, M. (2002). Depth-area-volume and hydroperiod relationships of ephemeral (vernal) forest pools in southern New England. Wetlands, 22(2), 247–255. 10.1672/0277-5212(2002)022[0247:DAVAHR]2.0.CO;2

9. Buxton, V. L., & Sperry, J. H. (2017). Reproductive decisions in anurans: a review of how predation and competition affects the deposition of eggs and tadpoles. BioScience, 67(1), 26–38. 10.1093/biosci/biw149

10. Cayuela, H., Valenzuela-Sánchez, A., Teulier, L., Martínez-Solano, Í., Léna, J. P., Merilä, J.,… & Schmidt, B. R. (2020). Determinants and consequences of dispersal in vertebrates with complex life cycles: a review of pond-breeding amphibians. The Quarterly Review of Biology, 95(1), 1–36. 10.1086/707862

11. Department of Agriculture, Government of Maharashtra (2023), Rainfall Recording and Analysis, https://maharain.maharashtra.gov.in/. Accessed on December 23, 2023.

12. Gaitonde, N., Giri, V., & Kunte, K. (2016). ‘On the rocks’: reproductive biology of the endemic toad Xanthophryne (Anura: Bufonidae) from the Western Ghats, India. Journal of Natural History, 50(39-40), 2557–2572. 10.1080/00222933.2016.1200686

13. Girish, K. G., & Krishnamurthy, S. V. B. (2009). Distribution of tadpoles of large wrinkled frog Nyctibatrachus major in central Western Ghats: influence of habitat variables. Acta Herpetologica, 4(2), 153–160. 10.13128/Acta_Herpetol-3417

14. Gosner, K. L. (1960). A simplified table for staging anuran embryos and larvae with notes on identification. Herpetologica, 16(3), 183–190. https://www.jstor.org/stable/3890061

15. Heyer, W. R., McDiarmid, R. W., & Weigmann, D. L. (1975). Tadpoles, predation and pond habitats in the tropics. Biotropica, 100–111. 10.2307/2989753

16. Jithin, V., Johnson, J. A., & Das, A. (2022). Influence of check dams on the activity pattern and morphometric traits of overwintering tadpoles in the Western Himalaya. Limnologica, 95, 125992. 10.1016/j.limno.2022.125992

17. Jithin, V., Rane, M., Watve, A., & Naniwadekar, R. (2023). Orchards and paddy differentially impact rock outcrop amphibians: Insights from community-and species-level responses. bioRxiv, 2023–10. 10.1101/2023.10.03.560737

18. Jocque, M., Vanschoenwinkel, B., & Brendonck, L. U. C. (2010). Freshwater rock pools: a review of habitat characteristics, faunal diversity and conservation value. Freshwater Biology, 55(8), 1587–1602. 10.1111/j.1365-2427.2010.02402.x

19. Kulkarni, A., Vijayan, S., Shigwan, B. K., Padhye, S., Dahanukar, N., & Datar, M. N. (2022). Environmental determinants of aquatic vegetation in rock pools of northern Western Ghats, India. Fundamental and Applied Limnology, 196(1), 15–26. 10.1127/fal/2022/1429

20. Kulkarni, M. R., Paripatyadar, S. V., Naniwadekar, R., & Joshi, J. (2023). Local-scale abiotic factors influence the organization of rock pool arthropod communities in a biodiversity hotspot. Limnology and Oceanography, 68(10), 2375–2388. 10.1002/lno.12427

21. Liedtke, H. C., Wiens, J. J., & Gomez-Mestre, I. (2022). The evolution of reproductive modes and life cycles in amphibians. Nature Communications, 13(1), 7039. 10.1038/s41467-022-34474-4

22. Luedtke, J. A., Chanson, J., Neam, K., Hobin, L., Maciel, A. O., Catenazzi, A., … & Stuart, S. N. (2023). Ongoing declines for the world’s amphibians in the face of emerging threats. Nature, 622(7982), 308–314. 10.1038/s41586-023-06578-4

23. Mogali, S. M., Shanbhag, B. A., & Saidapur, S. K. (2022). Predatory influence of dragonfly larvae and water scorpions on eggs and tadpoles of Indosylvirana temporalis (Anura: Ranidae). Phyllomedusa: Journal of Herpetology, 21(1), 51–57. 10.11606/issn.2316-9079.v21i1p51-57

24. Ovaskainen, O., & Abrego, N., (2020). Joint species distribution modelling: With applications in R. Cambridge University Press.

25. Ovaskainen, O., Tikhonov, G., Norberg, A., Guillaume Blanchet, F., Duan, L., Dunson, D., … & Abrego, N. (2017). How to make more out of community data? A conceptual framework and its implementation as models and software. Ecology letters, 20(5), 561–576. 10.1111/ele.12757

26. Priti, H., Gururaja, K. V., Aravind, N. A., & Ravikanth, G. (2021). Influence of microhabitat on the distribution of tadpoles of three endemic Nyctibatrachus species (Nyctibatrachidae) from the Western Ghats, India. Biotropica, 53(6), 1475–1485. 10.1111/btp.12988

27. R Core Team, 2024. R: A Language and Environment for Statistical Computing. R Foundation for Statistical Computing, Vienna, Austria.

28. Raj, P., Vasudevan, K., Kagarwal, R., Dutta, S. K., Sahoo, G., Mahapatra, S., … & Dubois, A. Larval morphology of selected anuran species from India. Alytes, 2023, 39–40: 1–140. https://www.biotaxa.org/Alytes/issue/view/10715/1141

29. Rothenberger, M. B., & Baranovic, A. (2021). Predator–prey relationships within natural, restored, and created vernal pools. Restoration Ecology, 29(1), e13308. 10.1111/rec.13308

30. Rothenberger, M. B., Vera, M. K., Germanoski, D., & Ramirez, E. (2019). Comparing amphibian habitat quality and functional success among natural, restored, and created vernal pools. Restoration ecology, 27(4), 881–891. 10.1111/rec.12922

31. Schiesari, L., Sgambatti Monteiro, A., Ilha, P., Pope, N., & Corrêa, D. T. (2018). The ecology of a system of natural mesocosms: Rock pools in the Atlantic Forest. Freshwater Biology, 63(9), 1077–1087. 10.1111/fwb.13118

32. Scott Jr, N. J., & Woodward, B. D. (1994). Surveys at breeding sites. In: Heyer et al. (Eds.) Measuring and monitoring biological diversity. Standard methods for amphibians. Smithsonian Books, Washington, D.C.

33. Sheth, S. D., Padhye, A. D., & Ghate, H. V. (2019). Factors affecting aquatic beetle communities of Northern Western Ghats of India (Arthropoda: Insecta: Coleoptera). Annales de Limnologie-International Journal of Limnology 55(1), 1–12. 10.1051/limn/2018030

34. Skelly, D. K. (1997). Tadpole communities: pond permanence and predation are powerful forces shaping the structure of tadpole communities. American Scientist, 85(1), 36–45. https://www.jstor.org/stable/27856689

35. Thomas, A., Das, S., & Manish, K. (2019). Influence of stream habitat variables on distribution and abundance of tadpoles of the endangered Purple frog, Nasikabatrachus sahyadrensis (Anura: Nasikabatrachidae). Journal of Asia-Pacific Biodiversity, 12(2), 144–151. 10.1016/j.japb.2019.01.009

36. Tikhonov, G., Opedal, Ø. H., Abrego, N., Lehikoinen, A., de Jonge, M. M., Oksanen, J., & Ovaskainen, O. (2020). Joint species distribution modelling with the R-package Hmsc. Methods in Ecology and Evolution, 11(3), 442–447. 10.1111/2041-210X.13345

37. Valdez, J. W. (2020). Arthropods as vertebrate predators: A review of global patterns. Global Ecology and Biogeography, 29(10), 1691–1703. 10.1111/geb.13157

38. Van Buskirk, J., & Smith, D. C. (2021). Ecological causes of fluctuating natural selection on habitat choice in an amphibian. Evolution, 75(7), 1862–1877. 10.1111/evo.14282

39. Vera Candioti, F., Baldo, D., Grosjean, S., Pereyra, M. O., & Nori, J. (2023). Global shortfalls of knowledge on anuran tadpoles. npj Biodiversity, 2(1), 22. 10.1038/s44185-023-00027-1

40. Watve, A. (2013). Status review of Rocky plateaus in the northern Western Ghats and Konkan region of Maharashtra, India with recommendations for conservation and management. Journal of Threatened taxa, 5(5), 3935–3962. 10.11609/JoTT.o3372.3935-62

